# South Asian Maternal Lineage haplogroup R30 Provides Phylogenetic Evidence of human dispersal across South Asia

**DOI:** 10.64898/2026.04.29.721543

**Authors:** Shailesh Desai, Vivek Adhikary, Mohak Bhattacharyya, Manoj Kumar Tharu, Abhishek Sharma, Jaison Jeevan Sequeira, Rudra kumar Pandey, Pratik Pandey, Shivanand S. Shendre, Alnoman Mundher Tayyeh, Sajitha Lulu S, Mohammed S. Mustak, Michael Petraglia, Gyaneshwer Chaubey

## Abstract

South Asia is central to debates on early human dispersals, particularly the Out of Africa model and Eurasian colonization. Studies of M haplogroups have been used to support both Northern and Southern route hypotheses, but current archaeological and genetic evidence in the region remains contradictory. In the present work, we find that in addition to haplogroup M lineages, a few R lineages exhibit ancient, locally rooted variation, with R30 being one of the widespread haplogroup of R lineages across South Asia. To better understand South Asian demographic history, we investigated the phylogeographic distribution of haplogroup R30, an indigenous lineage. We used 190 complete modern and ancient sequences from diverse mainland and island populations including incorporation of 44 newly generated sequences which enabled the refinement of the R30 phylogeny and the identification of a novel basal lineage, R30c. Bayesian and ρ-based age estimates suggest that R30 originated in the Indian subcontinent ~50 kya. Early diversification likely occurred in Northern India, giving rise to R30b (~44 kya), while R30a and R30c differentiated primarily in Southern India. Several subclades of haplogroup R30 exhibit strong signatures of founder effects, particularly among the language isolate Vedda of Sri Lanka, Uru Kurumban of Southern India, and the populations of the Lakshadweep archipelago. Bayesian skyline analyses indicate long-term demographic stability followed by rapid lineage expansion ~20 kya and more recent declines consistent with localised drift and relatively recent founder events. The presence of early-diverging R30 lineages in Thailand and Indonesia further supports long-term connections between South and Southeast Asia. Overall, archaeological and genetic evidence point towards the multiple migrations for South Asia colonizations.

## Introduction

Maternally inherited mitochondrial DNA (mtDNA) serves as a preferred marker for investigating lineage origins, human settlement, and colonization due to its rapid evolutionary rate and non-recombining uniparental inheritance. There is now growing evidence that the colonization of Eurasia by modern humans was not a simple process but consisted of multiple waves of Out of Africa (OOA) migration from the late middle Pleistocene to late Pleistocene (Groucutt et al., 2018; Hershkovitz et al., 2018; Freidline et al., 2023). A recent study suggested that an Out of Africa migration occurred ~70-60 kya at a large scale, but populations used the Persian Plateau as a hub, from where they then spread out to the rest of Eurasia (Vallini et al., 2024). Study of the descendants of this large-scale migration revealed that Eurasian haplogroups M and N represent the primary lineages that colonized Eurasia, with all contemporary humans tracing their ancestry to Africa. This Out of Africa migration led to the widespread colonization of Eurasia through Southern and Northern routes approximately 70-60 kya (Ingman et al., 2000; Maca-Meyer et al., 2001; Macaulay et al., 2005; Torroni et al., 2006; Cabrera et al., 2018; Vallini et al., 2024; Gandini et al., 2025). The existence of a Southern route is indicated by the presence of Middle Palaeolithic stone tool assemblages across the Arabian Peninsula (Petraglia and Alsharekh, 2003; Rose and Petraglia, 2009; Armitage et al., 2011; Tobler et al., 2023; Hallett et al., 2025) and genetic studies that reveal numerous indigenous M and N lineages (Kivisild et al., 2003; Macaulay et al., 2005; Merriwether et al., 2005; Thangaraj et al., 2006). Conversely, a Northern route through the Sinai Peninsula is supported by paleontological and archaeological evidence, including the discovery of anatomically modern human remains and stone artefacts in the Levant (Shea, 2003), and the considerable mean age of M and several N haplogroups in East and Southeast Asia (Larruga et al., 2017) alongside the absence of deep indigenous clades in the Arabian Peninsula (Fregel et al., 2015). However, the existence of indigenous M (Metspalu et al., 2004; Chandrasekar et al., 2009) and R (Palanichamy et al., 2004; Chaubey et al., 2008; Maji et al., 2008; Thangaraj et al., 2009) lineages, and their deep coalescence ages in India and Sri Lanka (Chandrasekar et al. 2009; Chaubey et al. 2008; Welikala et al., 2024), contradicts the Northern Route hypothesis. The presence of Middle Palaeolithic assemblages before MIS 5 and as recent as 30 kya (Petraglia et al., 2009; Petraglia et al., 2010; Akhilesh et al., 2018; Anil et al., 2022 & 2024), and the coalescence of South Asian lineages to Africa dating to no more than ~50 kya has raised important questions about the discordance between archaeological and genetic evidence, which still requires explanation.

Extensive research on maternal haplogroup M in relation to Out of Africa migrations has supported a Southern coastal route, specifically Andamanese-specific subclades such as M31 and M32 that support early coastal migrations from Africa around 70-60 kya (Thangaraj et al. 2005). For instance, investigations into haplogroup M have highlighted its deep coalescence ages (~55-50 kya in India) and widespread distribution across Asia and Australia, often interpreted as markers of the initial Southern dispersal. However, R lineages have been underrepresented in phylogeographic reconstructions despite their indigenous status in the region (Chandrasekar et al. 2009; Chaubey et al. 2008). These R lineages exhibit deep-rooted antiquity in the Indian subcontinent, with divergence dates for haplogroup R and its subclades often predating those of many M subclades, suggesting their potential as equally crucial markers for understanding early human migrations, regional demographic histories, and possible inland or alternative dispersal pathways (Palanichamy et al. 2004; Sun et al. 2006). This discrepancy in focus may stem from the higher diversity and frequency of M in relict populations like the Andamanese and Southeast Asian groups, yet emerging evidence from complete mtDNA sequencing indicates that autochthonous R subclades such as R5, R6, R7, R8, R30, and R31 arose in situ in South Asia, further underscoring the need for balanced inquiry into both macrohaplogroups (Palanichamy et al. 2004; Chandrasekar et al. 2009; Chaubey et al. 2008; Desai et al. 2026).

To provide phylogenetically resolved insights into Southern dispersal routes (Inland vs Coastal), here we focus on haplogroup R30, one of the most widespread indigenous South Asian lineages identified in diverse populations, including those from Northern and Southern India, Tibet, Sri Lanka, Thailand, and Indonesia. Recently this particular haplogroup has been recorded predominantly in island populations of Lakshadweep archipelago (Mustak et al., 2019; Tayyeh et al., 2023) as well as in the Vedda, aboriginal populations of Sri Lanka (Welikala et al., 2024). Despite the widespread occurrence of R30 in island populations, this haplogroup has been understudied. Hence, we analyze haplogroup R30’s detailed phylogeographic distribution across mainland and island groups using a comprehensive dataset of 190 modern and ancient complete sequences, including 44 newly generated ones.

## Methodology

### Sample collection and data generation

Samples were collected in accordance with guidelines and ethical permissions obtained from the Banaras Hindu University, Varanasi, India. Each individual was verbally informed and consent was obtained from them for use in biological studies. We collected 3-5 mL blood from each individual. Through the interview we ensured that they were from particular geographic areas for at least 3 generations, and not represented in our previously collected samples. In total, we collected 243 blood samples. Samples were stored and processed at the Gyan Lab, Cytogenetics Unit, Department of Zoology, Banaras Hindu University, Varanasi. Among these, 239 samples were collected from Amini island of Lakshadweep Archipelago, and 4 samples were collected from Tamilnadu state of Mainland India.

We amplified hyper variable segments, HVS-I (at 16024-16383) and HVS-II (at 57-372) of the mtDNA control region, using PCR. DNA stocks were diluted in TE buffer, from their respective concentrations to the standard working concentration of 10 ng/µl. Working primers (10µM) were made by diluting stock primers (100µM) in the TE buffer.

PCR was then performed by using 20µl total PCR mix, containing 2µl of working template, 0.5µl of each diluted primers (forward and reverse) and 10µl of EmeraldAmp® GT PCR Master Mix (Takara Bio Inc, Kusatsu, Japan), dissolved in 7µl of nuclease-free water. The amplification was performed using QIAamplifer 96 thermal cycler (QIAGEN GmbH, Hilden, Germany). Amplification was performed using set parameters as first denaturation at 95°C for 4 minutes, followed by 35 cycles of 30 seconds at 95°C (denaturation), 40 seconds at optimized annealing temperatures and 1 minute at 72°C (extension), further followed by final extension at 72°C for 7 minutes. All sets of reactions were run with a positive and negative control (as also outlined in Das et al., 2025). After quality control in 2% agarose gel electrophoresis, sequence data was obtained by performing standard Sanger sequencing procedure.

After successful sequencing, we manually checked the data and assigned the haplogroup. Among these 243 samples, 44 belong to R30, which was further processed with the same procedure, but for complete mtDNA sequence with approach outlined at our previous work (Desai et al., 2026).

### Data Curation and compilation

We sought the optimum process to screen the available data by investigating the available literature and the Mitomapper and yFull tree to see the potential for R30 haplogroup sequences. However, upon observing these two online tools, we found that the datasets were classifying the sequences of haplogroup B5 and haplogroup C4 into R30 due to some common mutations. We therefore manually assigned each variant to haplogroups, and we used MtPhyl for preparing the most parsimonious tree. However, since MtPhyl is not updated with the latest available haplogroup classification, we manually curated the rest of the trees. In addition, we manually reconstructed sequences of the Vedda (Welikala et al., 2024). For the above mentioned process and for further downstream analysis, we first aligned the data with rCRS references using the MUSCLE utility, available in MEGA v7.0, and aligned fasta files for further analysis. All R30 sequences and their source are available in Supplementary table 1. In total we generated 44 new sequences and 146 sequences were incorporated from other studies.

### Time calculations and Demographic analysis

The curated MtPhyl tree was used to prepare the RDF file in NETWORK for the calculation of Rho (ρ) and standard deviation (SD) values. The RDF file was manually prepared and visually inspected to minimize errors during data entry. The Rho values were then converted into time estimates using the formula provided by Soares et al. (2009). Previous research has highlighted the inconsistency in different studies for using different mutation rates as well as methods (Rho (ρ) and Beast) (Gandini et al., 2025), hence throughout this study, we have mainly used Rho (ρ) base methods for discussion. The aligned sequences were subsequently used to generate the XML file in BEAUti, a utility of Beast. Prior to this, the best-fit substitution model was selected using jModelTest2, and a mutation rate of 2.514 × 10^−8^ (Silva et al., 2017) was applied. The resulting XML files were analyzed in BEAST with 60 million iterations. All output parameters were assessed in Tracer to ensure effective sample size (ESS) values exceeded 200. Tracer was further used to generate the Bayesian skyline plot for the specific R30 haplogroup.

## Results and Discussion

### Refining the R30 haplogroup tree

The inclusion of our 44 novel samples, together with previously published R30 sequences, allowed us to revisit and refine the overall phylogeny of haplogroup R30. Haplogroup R30 was first reported by Palanichamy et al. (2004), subsequently refined by Chaubey et al. (2007), and more recently updated by Sylvester et al. (2019). R30 has been divided into two major subclades, R30a and R30b. The former is supported by ten coding-region substitutions, whereas the latter is defined by 24 mutations across both coding and control regions (Fig. 1). Several additional subclades within R30a and a few within R30b have been proposed; however, most were defined on the basis of limited sample sizes. Here, we define for the first time a basal branch of R30, designated R30c. This lineage is notable for comprising samples from Southern India and a single sample from Thailand. The R30c clade was further subdivided into R30c1 and R30c1a (Fig. 1 and Supplementary Figure 1).

**Fig 1.**
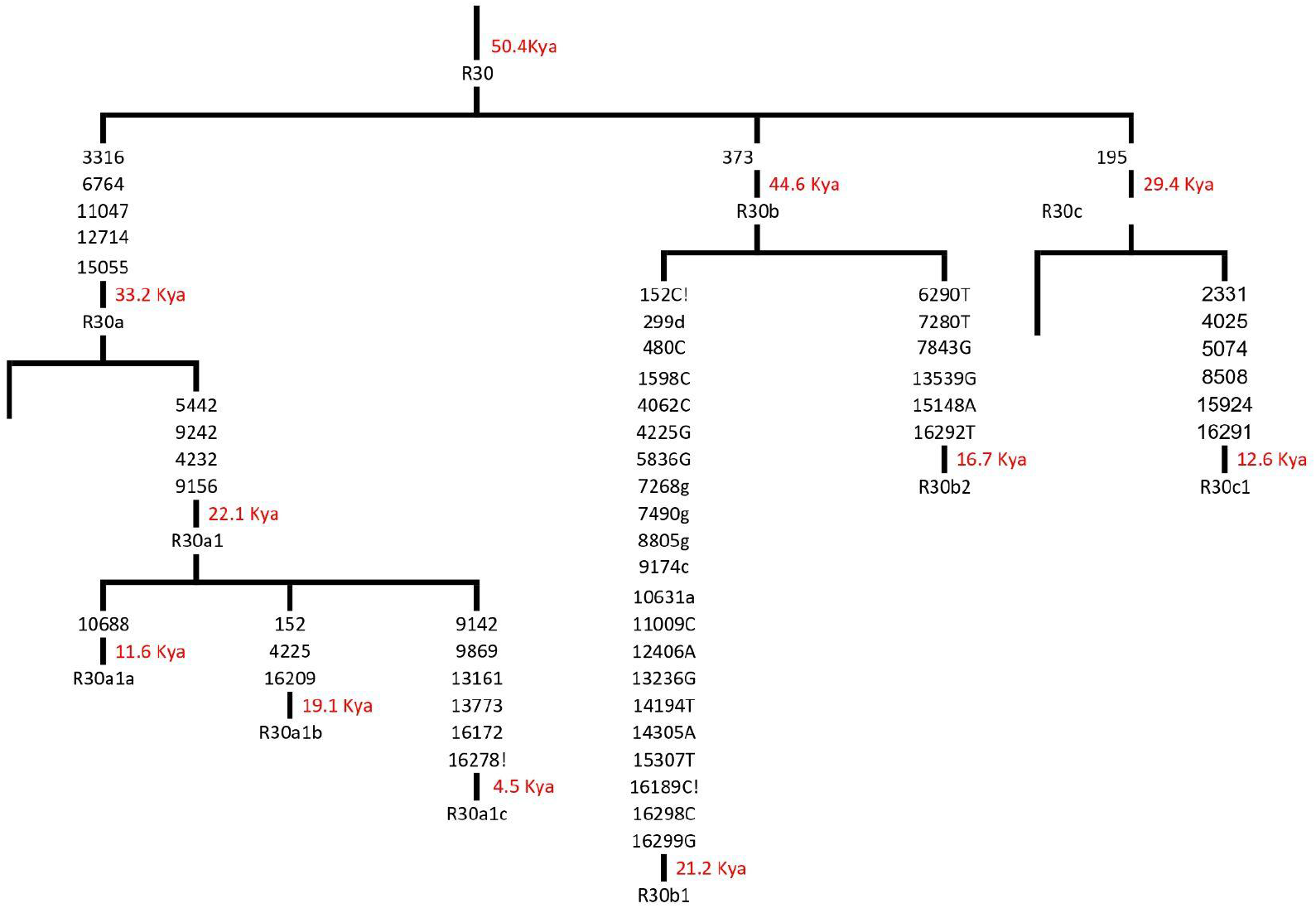
Refined R30 clade tree contains novel clades with time calculated from Rho base methods.

Among all R30 clades, R30a is the most extensively characterised, reflecting its widespread distribution in Indian populations. Here we further define two new sub-branches within this lineage: R30a1b1 and R30a1b1a. We also note that the recently described R30a1c1 (Sylvester et al., 2019) appears to be restricted to the Uru Kurumban population of Southern India, where it likely represents a strong founder lineage (Supplementary Figure 1).

### Demographic analysis of R30

To investigate the demographic history of haplogroup R30, we reconstructed a Bayesian skyline plot (BSP) using BEAST2. The BSP of R30 reveals a notable demographic pattern. The effective population size begins to increase shortly before 20 kya, continues rising until approximately 2–3 kya, and then shows a recent decline (Figure 2). Interestingly, the effective population size appears relatively stable prior to 20 kya, suggesting demographic stability during the earlier phase of its presence in the Indian subcontinent (approximately 50–20 kya). This can be explained by the relatively large number of mutations in several R30a and R30b subclades, the bearers of this lineage seem to have had demographic difficulties and remained as a stagnant population for a long time (Figure 1).

**Fig 2.**
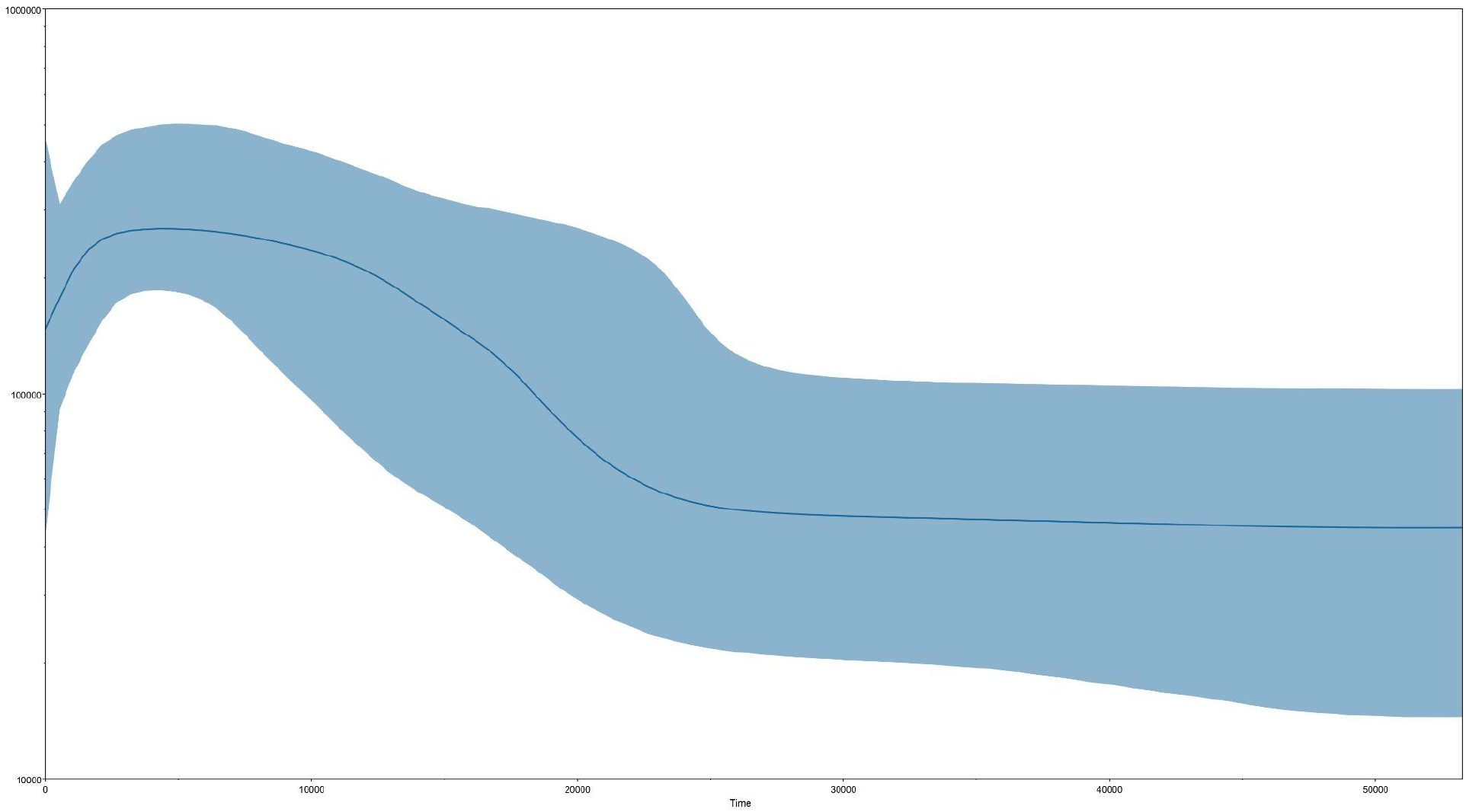
Bayesian skyline plot of R30 haplogroup contains all samples indicating the long term stable effective population size and increase at 20 kya.

The more recent decline in effective population size may reflect founder effects or increased endogamy within specific R30 subclades. For example, R30b2a2a1 shows evidence of a founder event in the Amini Islands, while R30b2a has been reported among the Vedda, population of Sri Lanka (Ranaweera et al., 2014; Chaubey, 2014; Welikala et al., 2024). Similarly, a strong founder effect has been documented in the Urali Kuruman population of

Southern India, where 87% of individuals carry the R30a1c1 lineage (Sylvester et al., 2019). Taken together, these multiple founder events across different subclades of R30 may explain the recent decline observed in the BSP. Notably, R30 may represent one of the few haplogroups demonstrating multiple independent founder events within distinct subclades.

### Spread of R30 into the Indian subcontinent

Our Bayesian and ρ-based age estimates indicate that haplogroup R30 originated in India prior to ~50 kya (Supplementary Fig 1; Figure 3). While recent whole-genome coalescent analyses suggest a demographic arrival in South Asia around ~45-50 kya (Kerdoncuff et al., 2025), uniparental markers often yield deeper temporal calibrations. Autosomal models, which rely on average population-level splits and identity-by-descent (IBD) sharing, can occasionally compress early evolutionary timelines due to extensive recombination and subsequent Holocene-era bottlenecks (Li and Durbin 2012). In contrast, the unbroken maternal line represented by haplogroup R30 provides a direct window into the earliest arrival of anatomically modern humans in the region. These data support a model where the initial entry and successful localised mutation of basal macrohaplogroups in South Asia predates the ~50 kya timeline, indicating a deep, continuous maternal heritage.

**Fig 3.**
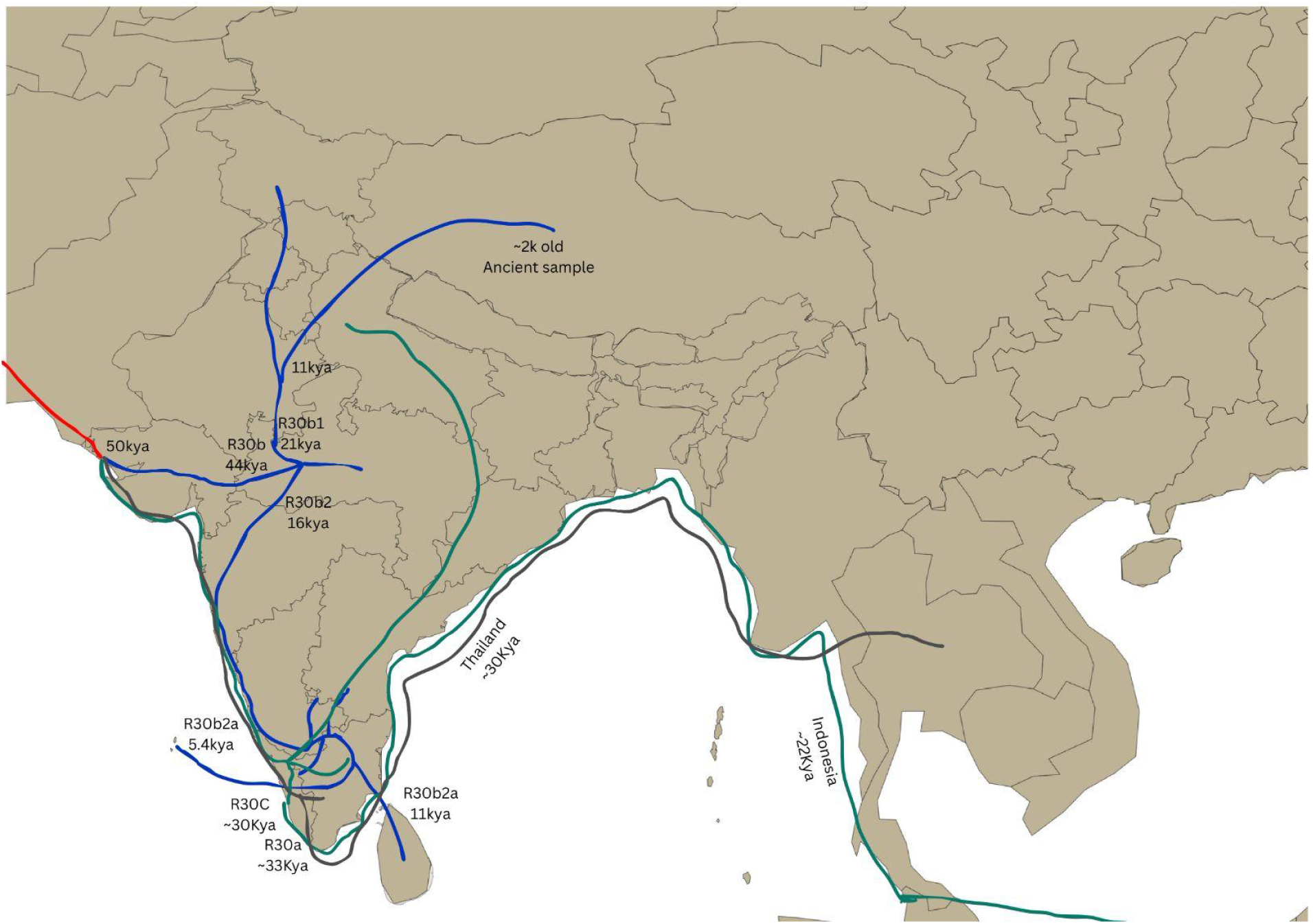
R30 haplogroup Distribution across the Indian subcontinent, where dark blue indicates R30b, Green indicates R30a, and Brown indicates the R30c.

The earliest differentiation appears to have occurred in Northern India, giving rise to haplogroup R30b at ~44 kya (Supplementary Fig 1). R30b subsequently diverged into R30b1 and R30b2 (Figure 3). At present, R30b1 is largely confined to Northern India, including the states of Punjab, Jammu and Kashmir, Uttar Pradesh, and Gujarat. R30b1 started diversifying within North India ~21.2 kya, and notably (Supplementary Fig 1), R30b1 has also been identified in an ancient individual from Tibet, which is consistent with our inference that this lineage was primarily distributed in Northern India. These findings highlight the extent to which present-day Indian populations may retain signals of ancient population structure, an important consideration given the limited availability of deep-time ancient DNA from the Indian subcontinent. Interestingly, in the Northern India, late Pleistocene Microlithic sites have been identified where R30b differentiated, noted by archaeological findings at Dhaba ~48 kya (Clarkson et al., 2020), Mehtakheri ~45 kya (Mishra et al., 2013), and Budha Pushkar ~28 kya (Blinkhom et al., 2018). Though this evidence does not necessarily indicate that these sites were occupied by R30 haplogroup-bearer groups, it does add validation to a human presence in this area with supporting archaeological and genetic information.

R30b2 diversified in Northern India and subsequently spread toward Southern regions around 16.7 kya (Supplementary Fig 1). This divergence timeline is notable as it corresponds to the environmental transitions following the Last Glacial Maximum (LGM, ~26–19 kya). During the height of the LGM, severe aridity and cooler temperatures across the subcontinent likely restricted human populations to localized ecological refugia. As the climate ameliorated and the South Asian monsoon systems strengthened during the post-LGM period, and previously inhospitable regions became viable for occupation, opening up new habitable corridors. Therefore, the simultaneous diversification and Southward expansion of R30b2 at ~16.7 kya appears directly coupled with paleoenvironmental shifts, indicating that post-glacial climatic changes facilitated both demographic expansions out of refugia and lineage differentiation. The subclade R30b2a has been reported at high frequency among the Vedda in Sri Lanka (Ranaweera et al., 2014; Chaubey, 2014; Welikala et al., 2024). Our age estimates suggest that this lineage entered Sri Lanka from India around ~11 kya and subsequently underwent a strong founder effect (Supplementary Fig 1). The period between 10 and 7 kya is characterised by significant sea-level rise and the loss of the land bridge between India and Sri Lanka (Hashimi et al., 1995; Loveson & Nigam, 2019; Dubey et al., 2023) which may have contributed to population isolation and the observed founder events. It is important to note that Microlithic sites in Sri Lanka have been dated back to ~48 kya, indicating that the Vedda possess a more ancient lineage (Welikala et al., 2024) compared to the R30b2a lineage. Thus, R30b2a shows merely the connection between India and Sri Lanka, which seem to suffer from founder events due to loss of land connections. Apart from this, founder events are evident in the Amini and Chetlat islands of the Lakshadweep archipelago. The lineages in these islands appear to have diverged around ~5.4 kya. This may suggest the origin of the first peoples of the Amini and Chetlat islands (Supplementary Fig 1).

Our results support the hypothesis that the Lakshadweep Islands were inhabited at least by ~6 kya, rather than being occupied only within the last couple of millennia (Figure 3). Currently the Lakshadweep Archipelago contains around 36 islands in the Southern Arabian Sea, though archaeological records of the early period are absent. Previous study has revealed that sex biased ancestry exists among these islands, as mostly maternally Southern India and paternal ancestry related to Northern India (Mustak et al., 2019). Currently there is no evidence of aboriginal people in the Lakshadweep Islands, but have been suggested various history of settlement (Forbes 1979; Rao 1986; Gupchup 1997; Tripati 1999; Saigal 2000; Vijaya Kumar 2006). Hence based on this, it has been concluded that there was a human population presence on these islands before at least 3500 years ago (Kumar et al., 2024), which we further stretch back to ~5.4 kya.

A newly defined basal clade within R30, designated R30c, shows a younger coalescent age (~29 kya) compared to R30a and R30b. This lineage subsequently differentiated into R30c1 around ~12.6 kya (Supplementary Fig 1). R30c is primarily restricted to Southern India, with the exception of a single sample from Thailand. This Thailand sample shares a key mutation with the South Indian R30c lineages and diverges at ~29 kya. We therefore infer that this lineage likely split early from its South Indian counterparts and migrated towards Thailand along the eastern coastal corridor during that period (Figure 3). The Thailand sample also shares mutation 3316, which is one of the defining mutations of R30a. This shared mutation may explain why the Bayesian phylogeny clusters this sample within the broader R30a group. Such discrepancies between Bayesian and parsimony-based trees are not unexpected and may arise due to differences in model assumptions and algorithm approaches (Supplementary Figure 2).

Another split out of India is observed within the R30a1 lineage, and it appears to have diverged around ~21 kya, where it is currently present in Indonesia (Supplementary Figure 1). Taken together, these observations suggest the possibility of continued or recurrent gene flow between South Asia and Southeast Asia since the last ~30 kya. Importantly, the inferred migrations dated to ~29 kya (Indonesia) and ~21 kya (Thailand) should not be interpreted as excluding earlier dispersal events. In fact, deeper and more basal branches of several haplogroups have already been documented in Southeast Asia and Oceania (Gandini et al., 2025).

Within this broader framework, the R30 haplogroup provides direct phylogenetic evidence of connections that are consistent with the hypothesized dispersals along the Southern route during MIS 3 of the Late Pleistocene (~60–50 kya). Notably, R30 represents the first maternal haplogroup offering such a direct genetic link between South Asia and Southeast Asia along these proposed migration routes.

### Human Colonization of South Asia

South Asia has always been a critical point for discussion of the human colonization of Eurasia owing to its unique geographic location and its high genetic, cultural and linguistic diversity (Petraglia & Allchin, 2007). Based on evidence from archaeology and genetics, scholars have proposed two contrasting models to explain when and how modern humans first appeared in the subcontinent (Petraglia et al., 2009; Haslam et al., 2010; Petraglia et al., 2010; Mellars et al., 2013; Mishra et al., 2013). The ‘late colonization’ model combined archaeological and genetic evidence to suggest that modern humans arrived in South Asia after ~55 kya, surmised to be associated with microlithic technology, with expansions eastward to reach Australia by at least 50 kya (Macaulay et al., 2005; Mellars, 2006; Fernandes et al., 2012; Mishra et al., 2013; Lewis et al., 2014; Silva et al., 2017). In contrast, the original formulation of the ‘early colonization’ model was based primarily on archaeological records, which argued for earlier phases of out of Africa migrations, with the entry of modern humans in South Asia during MIS 5, carrying Middle Palaeolithic technology (Petraglia et al., 2009; Haslam et al., 2010; Petraglia et al., 2010). In the late colonization model modern humans using Microlithic toolkits between 55-40 kyr would have potentially encountered and replaced archaic hominins, whereas in the early model, modern humans using Middle Palaeolithic toolkits may have encountered archaic ancestors prior to the Toba super-eruption of 74 kyr, using similar tool assemblages (see for example, Mellars et al. 2013, figure 1). In recent years, fossil, genetic and archaeological evidence has indicated that earlier dispersals out of Africa occurred as early as ~200 kyr, indicating multiple migrations of modern humans, while supporting a spread out of Africa ~70-60 kya, with a major expansion across Eurasia ~45 kya (Bae et al. 2017; Vallini et al. 2024).

Our extensive analysis of R30 haplogroup suggests its origin ~50 kya in the Northern part of India, followed by subsequent diversification. This finding is consistent with previous studies indicating that the majority of South Asian ancestry and haplogroups rarely predate ~60 kya (Thangaraj et al., 2006; Chaubey et al., 2008; Larruga et al., 2017; Silva et al., 2017). For example, recent work on Sri Lankan populations, in combination with South Indian datasets, estimates haplogroup R7 to ~56 kya (Welikala et al., in Rev). Furthermore, when founder samples are excluded and only representative samples of all branches of R30 are considered, the origin of R30 is pushed back to ~58 kya, still younger than 60 kya. These studies suggest that present-day South Asian populations are largely descendants of major Out-of-Africa migrations occurring ~60–50 kya, likely via the southern dispersal route. In the context of migration routes into South Asia, previous studies have favored a Northern route, citing the lack of deeply rooted, autochthonous N(xR) and M lineages in India, particularly in the Northern regions of India (Marrero et al., 2016; Larruga et al., 2017). Notably, the mean age of haplogroup R in South Asia has been estimated at ~47 ± 10.7 kya (Larruga et al., 2017).

Several indigenous R subclades, such as R6 and R31, along with certain clades of U and JT, have estimated ages of at least ~40 kya (Larruga et al., 2017). Our recent estimate of the coalescence age of haplogroup R7, based on a large dataset, places it at ~56 kya in South India and Sri Lanka (Welikala et al., in review). Building on this, here we demonstrate that haplogroup R30 ranges to ~50 kya, with an initial spread into Northern India, with subsequent differentiation into subclades R30b and R30c occurring in Southern India. Among these, R30c appears to be restricted to South India, providing support for the southern route. We also observe substantial variation in the coalescence times of R30 subclades (R30a: ~33.2 kya; R30b: ~44.6 kya; R30c: ~29.4 kya). This range of variation may reflect the effects of early genetic drift in South Asian populations, which could lead to an underestimation of the true ages of these lineages.

Large-scale genomic studies indicate that contemporary South Asian populations share their most recent common ancestor (TMRCA) with African populations at ~50 kya (Kerdoncuff et al., 2025). In contrast, archaeological records document the early emergence and long persistence of Middle Paleolithic technologies in South Asia from ~385 kyr to as late as ~38 kya (Petraglia et al., 2009; Akhilesh et al., 2018; Anil et al., 2022, 2024) (Figure 4). This temporal discrepancy raises important questions about population continuity and cultural transmission.

**Fig 4.**
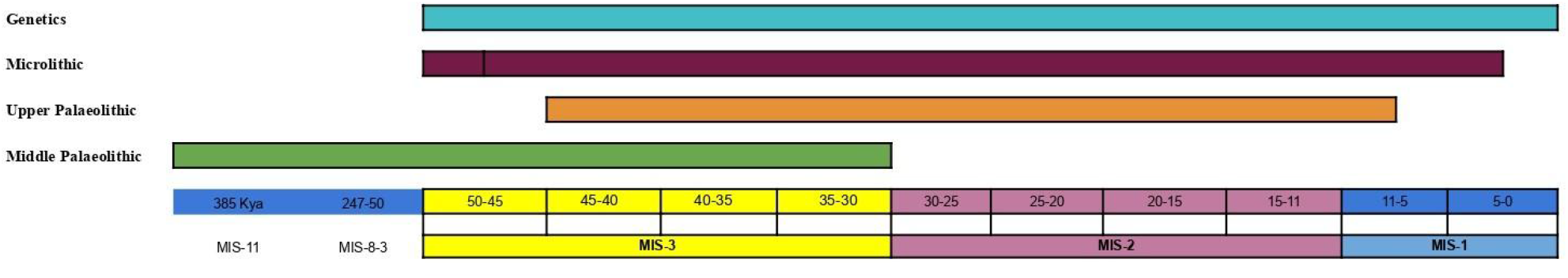
Graph showing the continuity of various tools, technique and genetics with temporal scale, in accordance with previously available data and present work.

Combination of genetic evidence with recent archaeological findings suggests the potential for two new scenarios to be considered in South Asia. One possibility is that endemic Middle Paleolithic assemblages were produced and used by archaic humans (possibly Neanderthals or Denisovans), and were later exchanged with modern humans with their genes, as indicated by introgression (Kerdoncuff et al., 2025). Indeed, such a scenario has been proposed in the Levant, where the sharing of Middle Paleolithic technology between Neanderthals and Homo sapiens has been suggested (Zaidner et al., 2025). Another scenario to consider involves evidence of admixture from ghost populations within South Asia, which may have used these tools and subsequently transmitted them to modern humans while contributing only limited genetic input (Mondal et al., 2016; 2018; 2019). Moreover, ancestry deconvolution of the inferred AASI (Ancient Ancestral South Indian) component reveals that it diverged from the Out-of-Africa source population earlier than present-day East Asian (Han) and Andamanese (Onge) populations (Yelmen et al., 2019). Importantly, a recent study (Kerdoncuff et al., 2025) reported that South Asian populations harbor a component of ancestry (~3%) that predates ~74 kya, marking the period of the Toba super-eruption, and aligning with earlier archaeological findings that indicate continuity both before and after this event (Petraglia et al., 2007). At present, both scenarios remain plausible, until clearer associations can be established between the use of Middle Paleolithic technology and other Homo groups in South Asia

In sum, South Asia represents a critical frontier for investigating the timing of Out-of-Africa migrations and provides an important context for examining the interplay between biological and cultural exchanges among human populations.

## Conclusion

South Asia serves as a critical crossroads for understanding the dispersal of anatomically modern humans from Africa. In the current study, we explored the phylogeographic dynamics of the indigenous maternal haplogroup R30 using 190 complete sequences. Our findings indicate that R30 originated in the Indian subcontinent prior to 50 kyr, representing a deep maternal foundation for early human colonisation. The topological divergence of R30 reveals a highly structured demographic history: R30b initially differentiated in Northern India and expanded Southward in response to post-LGM climatic improvements around 16.7 kya, while R30a and the newly identified basal clade R30c diversified primarily within Southern India. Notably, the presence of early-diverging R30 lineages in Thailand and Indonesia offers direct phylogenetic evidence of sustained human connectivity between South and Southeast Asia, following the proposed Southern route. Additionally, Bayesian demographic models illustrate a prolonged period of genetic stability, followed by a rapid population expansion around 20 kya. Subsequent localized declines correspond strongly with significant recent founder events seen in the Vedda of Sri Lanka, the Uru Kurumban of Southern India, and the isolated populations of the Lakshadweep archipelago. In sum, by integrating our uniparental data with recent whole-genome and archaeological findings, we have proposed scenarios that may accord well with the ~50 kya genomic threshold for the primary demographic expansion along with the deeper antiquity of regional tool cultures. Ultimately, the deep roots of haplogroup R30 highlight the intricate complexity of South Asian prehistory, calling for further high-resolution genomic investigations.

## Data Availability

The newly sequenced data will be submitted to the NCBI.

## Contributions

SD, MP, and GC conceived and designed the study. SD, VA, MB, MKT, AS, JJS, RKP, PP, SSS, AMT, SLS, MM and GC were involved in sample collection and performed the analyses. SD carried out the primary data processing and statistical analyses. SD, PP, MP, and GC wrote the original draft of the manuscript. All authors contributed to data interpretation, critically revised the manuscript for intellectual content, and approved the final version.

## Acknowledgments

S.D. is supported by the Council of Scientific and Industrial Research-Junior Research Fellowship (SRF-CSIR). G.C. is supported by Indian Council of Medical Research ad hoc grants (2021-6389 and 2021-11289) and Institute of Eminence, Banaras Hindu University (6031). R.K.P. is supported by the Indian Council of Medical Research-Senior Research Fellowship (2021-15707). The support and resources provided by the PARAM Shivay Facility under the National Supercomputing Mission, Government of India at the Indian Institute of Technology, Varanasi are gratefully acknowledged.

## Figure Legend

Supplementary Figure 1: Manually constructed Most parsimonious tree of R30s branches, and time calculation through Rho based methods.

Supplementary Figure 2. Bayesian analysis tree indicates the divergence among different samples and clades of R30a, R30b and R30c.

