## Supplementary table 1 and Supplementary Figure 2 for "South Asian Maternal Lineage haplogroup R30 Provides Phylogenetic Evidence of human dispersal across South Asia"

Supplementary Table 1: Details of each sample used in this study, their affiliation and sources.

| Sr.No | Accession Number | Population Name | Geographic Area (Country) | Source | Remark |
|---|---|---|---|---|---|
| 1 | DQ341064 | Africa | Africa | Torroni et al., 2006. | Outgroup |
| 2 | VE17 | Vedda | SriLanka | Welikala et al., 2024 | Reconstructed |
| 3 | VE15 | Vedda | SriLanka | Welikala et al., 2024 | Reconstructed |
| 4 | VE12 | Vedda | SriLanka | Welikala et al., 2024 | Reconstructed |
| 5 | VE09 | Vedda | SriLanka | Welikala et al., 2024 | Reconstructed |
| 6 | VE07 | Vedda | SriLanka | Welikala et al., 2024 | Reconstructed |
| 7 | VE06 | Vedda | SriLanka | Welikala et al., 2024 | Reconstructed |
| 8 | VE05 | Vedda | SriLanka | Welikala et al., 2024 | Reconstructed |
| 9 | VE02 | Vedda | SriLanka | Welikala et al., 2024 | Reconstructed |
| 10 | V3 | Vedda | SriLanka | Welikala et al., 2024 | Reconstructed |
| 11 | OQ707189 | Tibet | Tibet | Zhang et al., 2023 | Ancient (~2000 year old) |
| 12 | OP004763 | Gujarat | West part of India | Alqaisi et al., 2024; Desai et al., 2025 | |
| 13 | OP004762 | Gujarat | West part of India | Alqaisi et al., 2024; Desai et al., 2025 | |

| | | | | | |
|---|---|---|---|---|---|
| 14 | OP004761 | Gujarat | West part of India | Alqaisi et al., 2024; Desai et al., 2025 | |
| 15 | OM489685 | - | Sri Lanka | Direct Submitted | |
| 16 | NA21115 | Gujarat | West part of India | 1000 Genome Consortium | |
| 17 | NA21110 | Gujarat | West part of India | 1000 Genome Consortium | |
| 18 | NA21105 | Gujarat | West part of India | 1000 Genome Consortium | |
| 19 | NA20902 | Gujarat | West part of India | 1000 Genome Consortium | |
| 20 | NA20856 | Gujarat | West part of India | 1000 Genome Consortium | |
| 21 | MZ895060 | Vedda | Sri Lanka | Direct Submitted | Fernando and team |
| 22 | MT506310 | Parsi | India | Morawala-Patell et al., 2021 | |
| 23 | MN595720 | Pakistan | Pakistan | Rahman et al., 2021 | |
| 24 | MH844546 | Vedda | Sri lanka | Direct submission | Ranasinghe And team |
| 25 | MH830130 | Chetlet Island, Lakshadweep archipelago | India | Direct Submission | |
| 26 | MH830119 | Chetlet Island, Lakshadweep archipelago | India | Direct Submission | |

| | | | | |
|---|---|---|---|---|
| 27 | MH830118 | Chetlet Island, Lakshadweep archipelago | India | Direct Submission |
| 28 | MH830117 | Chetlet Island, Lakshadweep archipelago | India | Direct Submission |
| 29 | MH830116 | Chetlet Island, Lakshadweep archipelago | India | Direct Submission |
| 30 | MH830115 | Chetlet Island, Lakshadweep archipelago | India | Direct Submission |
| 31 | MH830114 | Chetlet Island, Lakshadweep archipelago | India | Direct Submission |
| 32 | MH830113 | Chetlet Island, Lakshadweep archipelago | India | Direct Submission |
| 33 | MH830112 | Chetlet Island, Lakshadweep archipelago | India | Direct Submission |
| 34 | MH830111 | Chetlet Island, Lakshadweep archipelago | India | Direct Submission |
| 35 | MH830110 | Chetlet Island, Lakshadweep archipelago | India | Direct Submission |
| 36 | MH830109 | Chetlet Island, Lakshadweep archipelago | India | Direct Submission |
| 37 | MH830108 | Chetlet | India | Direct |

| | | | | |
|---|---|---|---|---|
| | | Island, Lakshadweep archipelago | | Submission |
| 38 | MH830107 | Chetlet Island, Lakshadweep archipelago | India | Direct Submission |
| 39 | MH830106 | Chetlet Island, Lakshadweep archipelago | India | Direct Submission |
| 40 | MH830105 | Chetlet Island, Lakshadweep archipelago | India | Direct Submission |
| 41 | MH830103 | Chetlet Island, Lakshadweep archipelago | India | Direct Submission |
| 42 | MH830102 | Chetlet Island, Lakshadweep archipelago | India | Direct Submission |
| 43 | MH830101 | Chetlet Island, Lakshadweep archipelago | India | Direct Submission |
| 44 | MH830100 | Chetlet Island, Lakshadweep archipelago | India | Direct Submission |
| 45 | MH830099 | Chetlet Island, Lakshadweep archipelago | India | Direct Submission |
| 46 | MH830098 | Chetlet Island, Lakshadweep archipelago | India | Direct Submission |
| 47 | MF437223 | UAE | UAE | Aljasmi et al., 2020 |

| | | | | |
|---|---|---|---|---|
| 48 | MF437122 | UAE | UAE | Aljasmi et al., 2020 |
| 49 | LK-F04 | Amini Island, Lakshadweep archipelago | India | This Study |
| 50 | LK-E09 | Amini Island, Lakshadweep archipelago | India | This Study |
| 51 | LK-E04 | Amini Island, Lakshadweep archipelago | India | This Study |
| 52 | LK-E02 | Amini Island, Lakshadweep archipelago | India | This Study |
| 53 | LK-D09 | Amini Island, Lakshadweep archipelago | India | This Study |
| 54 | LK-D08 | Amini Island, Lakshadweep archipelago | India | This Study |
| 55 | LK-D05 | Amini Island, Lakshadweep archipelago | India | This Study |
| 56 | LK-D04 | Amini Island, Lakshadweep archipelago | India | This Study |
| 57 | LK-D03 | Amini Island, Lakshadweep archipelago | India | This Study |
| 58 | LK-D02 | Amini Island, Lakshadweep archipelago | India | This Study |

| | | | | |
|---|---|---|---|---|
| 59 | LK-C12 | Amini Island, Lakshadweep archipelago | India | This Study |
| 60 | LK-C10 | Amini Island, Lakshadweep archipelago | India | This Study |
| 61 | LK-C09 | Amini Island, Lakshadweep archipelago | India | This Study |
| 62 | LK-C08 | Amini Island, Lakshadweep archipelago | India | This Study |
| 63 | LK-C04 | Amini Island, Lakshadweep archipelago | India | This Study |
| 64 | LK-C03 | Amini Island, Lakshadweep archipelago | India | This Study |
| 65 | LK-C02 | Amini Island, Lakshadweep archipelago | India | This Study |
| 66 | LK-B12 | Amini Island, Lakshadweep archipelago | India | This Study |
| 67 | LK-B11 | Amini Island, Lakshadweep archipelago | India | This Study |
| 68 | LK-B10 | Amini Island, Lakshadweep archipelago | India | This Study |

| | | | | |
|---|---|---|---|---|
| 69 | LK-B09 | Amini Island, Lakshadweep archipelago | India | This Study |
| 70 | LK-B08 | Amini Island, Lakshadweep archipelago | India | This Study |
| 71 | LK-B05 | Amini Island, Lakshadweep archipelago | India | This Study |
| 72 | LK-B04 | Amini Island, Lakshadweep archipelago | India | This Study |
| 73 | LK-B02 | Amini Island, Lakshadweep archipelago | India | This Study |
| 74 | LK-B01 | Amini Island, Lakshadweep archipelago | India | This Study |
| 75 | LK-A11 | Amini Island, Lakshadweep archipelago | India | This Study |
| 76 | LK-A07 | Amini Island, Lakshadweep archipelago | India | This Study |
| 77 | LK-A06 | Amini Island, Lakshadweep archipelago | India | This Study |
| 78 | LK-A05 | Amini Island, Lakshadweep archipelago | India | This Study |
| 79 | LK-A03 | Amini Island, | India | This Study |

| | | | | |
|---|---|---|---|---|
| | | Lakshadweep archipelago | | |
| 80 | LK-A02 | Amini Island, Lakshadweep archipelago | India | This Study |
| 81 | LK-A01 | Amini Island, Lakshadweep archipelago | India | This Study |
| 82 | KY686208 | Indonesia | Indonesia | Silva et al., 2017 |
| 83 | KX467277 | Jammu and Kashmir | India | Sharma et al., 2018 |
| 84 | KX457111 | Mon | Thailand | Kutanan et al., 2017 |
| 85 | KU683258 | Uyghur | Xingjiang, China | Zheng et al., 2017 |
| 86 | KU683250 | Uyghur | Xingjiang, China | Zheng et al., 2017 |
| 87 | KF038316 | Surat, Gujarat | India | Direct Submission |
| 88 | KC911618 | Iran | Iran | Derenko et al., 2013 |
| 89 | KC911324 | Iran | Iran | Derenko et al., 2013 |
| 90 | JX462737 | South India | India | Khan et al., 2013 |
| 91 | JX462715 | South India | India | Khan et al., 2013 |
| 92 | JX462707 | South India | India | Khan et al., 2013 |
| 93 | JX462702 | South India | India | Khan et al., 2017 |

| | | | | |
|---|---|---|---|---|
| 94 | HG04018 | Indian Tamil | India | The 1000 Genomes Project Consortium, 2010 |
| 95 | HG03949 | Srilankan Tamil | Sri Lanka | The 1000 Genomes Project Consortium, 2010 |
| 96 | HG03937 | Bengali | Bangladesh | The 1000 Genomes Project Consortium, 2010 |
| 97 | HG03911 | Bengali | Bangladesh | The 1000 Genomes Project Consortium, 2010 |
| 98 | HG03887 | Srilankan Tamil | Sri Lanka | The 1000 Genomes Project Consortium, 2010 |
| 99 | HG03874 | Indian Tamil | India | The 1000 Genomes Project Consortium, 2010 |
| 100 | HG03872 | Indian Tamil | India | The 1000 Genomes Project Consortium, 2010 |

| | | | | |
|---|---|---|---|---|
| 101 | HG03788 | Indian Tamil | India | The 1000 Genomes Project Consortium, 2010 |
| 102 | HG03714 | Indian Tamil | India | The 1000 Genomes Project Consortium, 2010 |
| 103 | HG03680 | Srilankan Tamil | Sri Lanka | The 1000 Genomes Project Consortium, 2010 |
| 104 | HG03679 | Srilankan Tamil | Sri Lanka | The 1000 Genomes Project Consortium, 2010 |
| 105 | HG03645 | Srilankan Tamil | Sri Lanka | The 1000 Genomes Project Consortium, 2010 |
| 106 | HG02697 | Punjabi | Pakistan | The 1000 Genomes Project Consortium, 2010 |
| 107 | HG01593 | Punjabi | Pakistan | The 1000 Genomes Project Consortium, 2010 |

| | | | | |
|---|---|---|---|---|
| 108 | GU170818 | TamilNadu | India | Rani et al., 2010 |
| 109 | GU170815 | TamilNadu | India | Rani et al., 2010 |
| 110 | GQ337679 | India | India | Direct Submission |
| 111 | GQ337672 | Andhra Pradesh | India | Direct Submission |
| 112 | GQ337671 | India | India | Direct Submission |
| 113 | GQ337669 | India | India | Direct Submission |
| 114 | GQ337541 | India | India | Direct Submission |
| 115 | GA001422 | Indian Tamil | India | GenomeAsia 100K Consortium, 2019 |
| 116 | GA001409 | Indian Tamil | India | GenomeAsia 100K Consortium, 2019 |
| 117 | GA001399 | Indian Tamil | India | GenomeAsia 100K Consortium, 2019 |
| 118 | GA001398 | Srilankan Tamil | Srilanka | GenomeAsia 100K Consortium, 2019 |
| 119 | GA001312 | Urban Banglore | India | GenomeAsia 100K Consortium, 2019 |
| 120 | GA001296 | Urban Banglore | India | GenomeAsia 100K |

| | | | | |
|---|---|---|---|---|
| | | | | Consortium, 2019 |
| 121 | GA001265 | Urban Chennai | India | GenomeAsia 100K Consortium, 2019 |
| 122 | GA001253 | Urban Chennai | India | GenomeAsia 100K Consortium, 2019 |
| 123 | GA001196 | Mahar, North India | India | GenomeAsia 100K Consortium, 2019 |
| 124 | GA001157 | Paniya, South India | India | GenomeAsia 100K Consortium, 2019 |
| 125 | GA001149 | Paniya, South India | India | GenomeAsia 100K Consortium, 2019 |
| 126 | GA001148 | Paniya, South India | India | GenomeAsia 100K Consortium, 2019 |
| 127 | GA001144 | Lambada, South India | India | GenomeAsia 100K Consortium, 2019 |
| 128 | GA001142 | Lambada, South India | India | GenomeAsia 100K Consortium, 2019 |
| 129 | GA001101 | Iyer, South India | India | GenomeAsia 100K |

| | | | | | |
|---|---|---|---|---|---|
| | | | | Consortium, 2019 | |
| 130 | GA000965 | Chamar, North India | India | GenomeAsia 100K Consortium, 2019 | |
| 131 | FJ770961 | Tharu | Nepal | Fornarino et al., 2009 | |
| 132 | FJ004827 | Punjab | India | Chaubey et al., 2008 | |
| 133 | FJ004824 | Sinhalese | SriLanka | Chaubey et al., 2008 | |
| 134 | FJ004807 | Maharashtra | India | Chaubey et al., 2008 | |
| 135 | EF556148 | Indian jews | India | Behar et al., 2008 | |
| 136 | AY714050 | UttarPradesh | India | Palanichamy et al., 2004 | |
| 137 | AY714047 | UttarPradesh | India | Palanichamy et al., 2004 | |
| 138 | AY714032 | UttarPradesh | India | Palanichamy et al., 2004 | |
| 139 | AY714006 | Andhra Pradesh | India | Palanichamy et al., 2004 | |
| 140 | AY714001 | Andhra Pradesh | India | Palanichamy et al., 2004 | |
| 141 | AV5 | Vedda | Sri Lanka | Welikala et al., 2024 | Reconstructed |

| | | | | | |
|---|---|---|---|---|---|
| 142 | AV4 | Vedda | Sri Lanka | Welikala et al., 2024 | Reconstructed |
| 143 | AV3 | Vedda | Sri Lanka | Welikala et al., 2024 | Reconstructed |
| 144 | AV2 | Vedda | Sri Lanka | Welikala et al., 2024 | Reconstructed |
| 145 | 150LK | Amini Island, Lakshadweep archipelago | India | This Study | |
| 146 | 146LK | Amini Island, Lakshadweep archipelago | India | This Study | |
| 147 | 145LK | Amini Island, Lakshadweep archipelago | India | This Study | |
| 148 | 144LK | Amini Island, Lakshadweep archipelago | India | This Study | |
| 149 | 143LK | Amini Island, Lakshadweep archipelago | India | This Study | |
| 150 | 119LK | Amini Island, Lakshadweep archipelago | India | This Study | |
| 151 | 111LK | Amini Island, Lakshadweep archipelago | India | This Study | |
| 152 | GC_151 | | India | This Study | |
| 153 | GC_262 | | India | This Study | |
| 154 | T18 | | India | This Study | |
| 155 | T33 | | India | This Study | |

| | | | | |
|---|---|---|---|---|
| 156 | MH499962 | Urali Kurumans | India | Sylvester et al., 2019 |
| 157 | MH499963 | Urali Kurumans | India | Sylvester et al., 2019 |
| 158 | MH499964 | Urali Kurumans | India | Sylvester et al., 2019 |
| 159 | MH499965 | Urali Kurumans | India | Sylvester et al., 2019 |
| 160 | MH499966 | Urali Kurumans | India | Sylvester et al., 2019 |
| 161 | MH499967 | Urali Kurumans | India | Sylvester et al., 2019 |
| 162 | MH499968 | Urali Kurumans | India | Sylvester et al., 2019 |
| 163 | MH499969 | Urali Kurumans | India | Sylvester et al., 2019 |
| 164 | MH499970 | Urali Kurumans | India | Sylvester et al., 2019 |
| 165 | MH499971 | Urali Kurumans | India | Sylvester et al., 2019 |

| | | | | |
|---|---|---|---|---|
| 166 | MH499972 | Urali Kurumans | India | Sylvester et al., 2019 |
| 167 | MH499973 | Urali Kurumans | India | Sylvester et al., 2019 |
| 168 | MH499974 | Urali Kurumans | India | Sylvester et al., 2019 |
| 169 | MH499975 | Urali Kurumans | India | Sylvester et al., 2019 |
| 170 | MH499976 | Urali Kurumans | India | Sylvester et al., 2019 |
| 171 | MH499977 | Urali Kurumans | India | Sylvester et al., 2019 |
| 172 | MH499978 | Urali Kurumans | India | Sylvester et al., 2019 |
| 173 | MH499979 | Urali Kurumans | India | Sylvester et al., 2019 |
| 174 | MH499980 | Urali Kurumans | India | Sylvester et al., 2019 |
| 175 | MH499981 | Urali Kurumans | India | Sylvester et al., 2019 |

| | | | | |
|---|---|---|---|---|
| 176 | MH499982 | Urali Kurumans | India | Sylvester et al., 2019 |
| 177 | MH499983 | Urali Kurumans | India | Sylvester et al., 2019 |
| 178 | MH499984 | Urali Kurumans | India | Sylvester et al., 2019 |
| 179 | MH499985 | Urali Kurumans | India | Sylvester et al., 2019 |
| 180 | MH499986 | Urali Kurumans | India | Sylvester et al., 2019 |
| 181 | MH499987 | Urali Kurumans | India | Sylvester et al., 2019 |
| 182 | MH499988 | Urali Kurumans | India | Sylvester et al., 2019 |
| 183 | MH499989 | Urali Kurumans | India | Sylvester et al., 2019 |
| 184 | MH499990 | Urali Kurumans | India | Sylvester et al., 2019 |
| 185 | MH499991 | Urali Kurumans | India | Sylvester et al., 2019 |

| | | | | |
|---|---|---|---|---|
| 186 | MH499992 | Urali Kurumans | India | Sylvester et al., 2019 |
| 187 | MH499993 | Urali Kurumans | India | Sylvester et al., 2019 |
| 188 | MH499994 | Urali Kurumans | India | Sylvester et al., 2019 |
| 189 | MH499995 | Urali Kurumans | India | Sylvester et al., 2019 |
| 190 | MH499996 | Urali Kurumans | India | Sylvester et al., 2019 |

Supplementary Table 2: Frequency of different haplogroup in Amini populations (this was based on Control region sequences).

| Haplogroup | Number of Sample | Frequency |
|---|---|---|
| A | 5 | 0.02183406114 |
| HV | 15 | 0.06550218341 |
| M2 | 12 | 0.05240174672 |
| M3 | 17 | 0.07423580786 |
| M5 | 9 | 0.03930131004 |
| M18 | 7 | 0.03056768559 |
| M30 | 16 | 0.06986899563 |
| M33 | 25 | 0.1091703057 |
| M35 | 30 | 0.1310043668 |
| M64 | 6 | 0.02620087336 |
| M4'67 | 11 | 0.0480349345 |

| | | |
|---|---|---|
| R30 | 54 | 0.2358078603 |
| U2 | 7 | 0.03056768559 |
| U7 | 15 | 0.06550218341 |

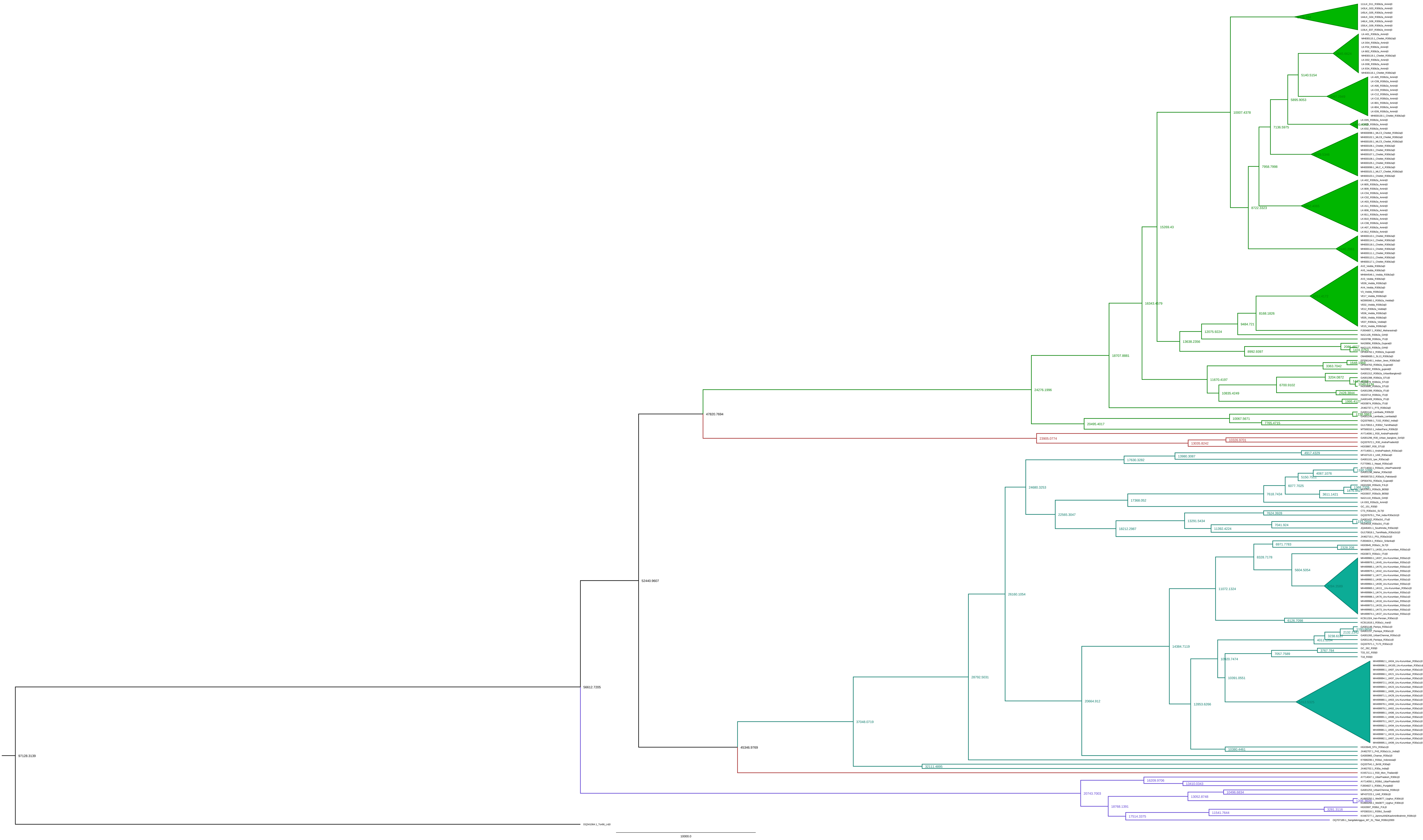
